# Systematic effects of retinotopic biases and category selectivity across human occipitotemporal cortex

**DOI:** 10.1101/2024.11.04.621840

**Authors:** Edward H Silson, Iris I A Groen, Chris I Baker

**Affiliations:** Department of Psychology, University of Edinburgh, Edinburgh, UK; Section on Learning and Plasticity, Laboratory of Brain and Cognition, National Institute of Mental Health, National Institutes of Health, Bethesda, USA; Institute for Informatics, University of Amsterdam, Amsterdam, The Netherlands

**Author notes:** Equal contribution (Edward H Silson, Iris I A Groen).

## Abstract

The organization of human visual cortex has traditionally been studied using two different methods: retinotopic mapping and category-selectivity mapping. Retinotopic mapping has identified a large number of systematic maps of the visual field, while category-selectivity mapping has identified clusters of neural populations that reliably respond more strongly to specific image categories such as objects, faces, scenes and body parts compared to other categories. While early investigations seemed to suggest that these two organizing principles were largely separated in the brain, with retinotopic maps in posterior visual cortex and category-selective regions in anterior visual cortex, recent work shows that category-selective regions overlap with retinotopic maps, giving rise to spatial visual field biases within these regions.

Here, we collected fMRI responses whilst performing both retinotopic and category mapping within the same participants, allowing detailed comparison of neural tuning for space and category at the single voxel level. We use these data to evaluate two previous proposals of how retinotopic biases relate to category-selectivity: 1) complementary quadrant biases (upper vs. lower contralateral visual field) inherited from early visual cortex explain the presence of paired regions selective for the *same category* across lateral and ventral occipitotemporal cortex (lOTC, vOTC); and 2) eccentricity biases (center vs. periphery of the visual field) explain the presence of selectivity for *different categories*, specifically differentiating face-versus scene-selectivity within the ventral surface.

Confirming and extending previous findings for a comprehensive set of face-, scene-, object, and place-selective regions of interest, we provide robust evidence that category-selective regions do not sample visual space uniformly, exhibiting systematic biases towards either the upper or lower field (all category regions) and center vs. periphery (face vs. place regions). Consistent with 1), we find that quadrant biases differ systematically between lateral and ventral OTC, with lateral regions showing systematic lower field biases and ventral regions showing upper field biases, differentiating regions selective for the same category in terms of their spatial bias. However, contrary to 2), we find that eccentricity tuning does not strongly predict the strength of face-or scene category-selectivity in a given voxel. Specifically, highly face-selective voxels are not solely confined to the fovea, and while most scene-selective voxels show peripheral tuning, highly scene-selective voxels actually show strong foveal tuning, particularly in anterior medial-ventral cortex.

Collectively, these results demonstrate that spatial biases in category-selective cortex are widespread and robust, whilst also suggesting there is no simple relation between spatial tuning and category-selectivity.

## Introduction

A hallmark of human occipitotemporal cortex organization is the presence of category-selective regions for faces, scenes, objects and bodies (Downing et al., 2001; Epstein & Kanwisher, 1998; Kanwisher et al., 1997; Kanwisher & Dilks, 2013; Malach et al., 2002) that occur at similar relative anatomical locations across individuals (Hasson et al., 2002; Levy et al., 2001), even in infancy (Deen et al., 2017). Together, these regions form an anatomically separated yet repeated functional organization across the lateral and ventral cortical surfaces, with corresponding selective regions in lateral (lOTC) and ventral (vOTC) occipitotemporal cortex **(Figure 1)**.

**Figure 1.**
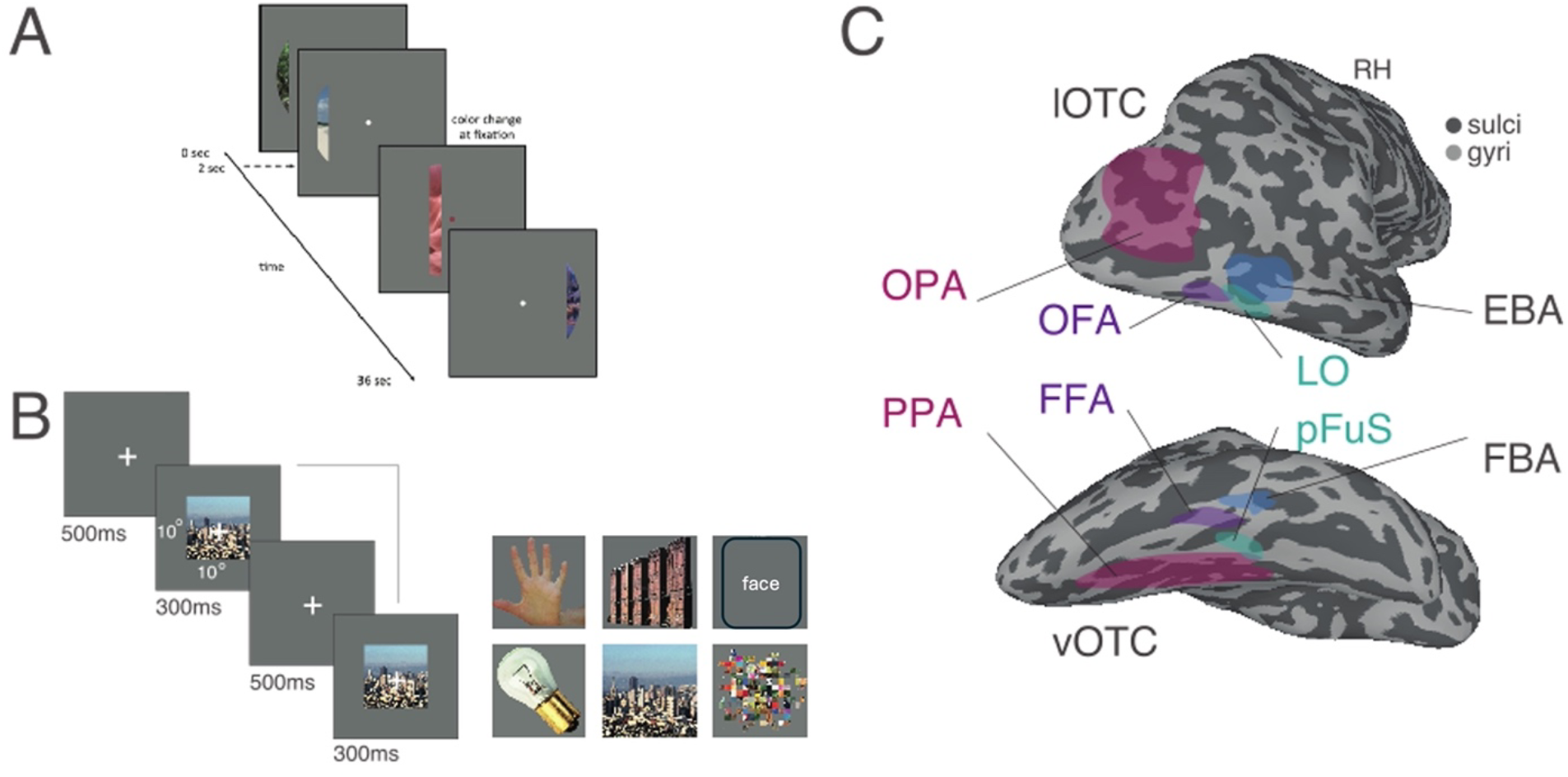
Task schematics and ROIs. **A**, Schematic of pRF mapping stimulus. A bar aperture traversed gradually through the visual field whilst revealing randomly selected scene-fragments. Participants fixated centrally and responded to a color-change at fixation via an MRI compatible response box. Bar apertures were taken from a circular aperture (diameter 20°). **B**, Schematic of category-localizer task. Stimuli from six different categories (bodies, buildings, faces, objects, scenes & scrambled) were presented at fixation in blocks. Participants fixated centrally and performed a 1-back task, responding via button press whenever the same image appeared sequentially. **C**, Group-level ROIs in the right hemisphere. Lateral (top) and ventral (bottom) partially inflated views of the right hemisphere are shown (sulci=dark gray, gyri=light gray). Overlaid in false color are the group-level ROIs: Scene-selective regions OPA and PPA (pink), Face-selective regions OFA and FFA (purple), Object-selective regions LO and pFuS (green) and body-selective regions EBA and FBA (blue). In each category the lateral and ventral pairs are spatially separated with one region on the lateral surface (OPA, OFA, LO & EBA) and one on the ventral surface (PPA, FFA, pFuS & FBA). Note that the lOTC ROIs are omitted from the vOTC view for simplicity.

A major question is what underlying principles drive this hallmark organization - why are these category regions located in these specific locations in cortex, and what determines their selectivity? Many possible explanations have been put forward, including a division of labor between early and late stages of visual processing (Taylor & Downing, 2011; Tsantani et al., 2021), anatomical or functional connections with downstream regions (Li et al., 2020; Saygin et al., 2016; Stevens et al., 2015), and representations of real-word size and animacy (Konkle & Caramazza, 2013; Konkle & Oliva, 2012). Another prominent explanation often invokes the relationship between retinotopic sampling and category selectivity (Groen et al., 2021; Hasson et al., 2002; Konkle & Caramazza, 2013; Konkle & Oliva, 2012; Levy et al., 2001; Op de Beeck et al., 2019; Silson et al., 2015; Taylor & Downing, 2011). Collectively these theories implicate retinotopic organization, such as large-scale eccentricity biases (Hasson et al., 2002; Levy et al., 2001) and division into dorsal and ventral maps, as retinotopic scaffolds that shape the formation of category regions in higher visual processing streams (Groen et al., 2021; Arcaro & Livingstone, 2021).

However, much is still unclear about the precise relationship between retinotopic biases and category-selectivity. Here, we sought to provide a comprehensive examination of the relation between visual field biases present in category-selective regions throughout OTC (n=30). Using separate fMRI measurements of visual field coverage and category-selectivity, we calculated the visual field biases exhibited by four pairs of category-selective regions spanning lOTC and vOTC, comprising scene-(Occipital Place Area OPA, Parahippoccampal Place Area, PPA), face-(Occipital Face Area, OFA, Fusiform Face Area, FFA), object-(Lateral Occipital Cortex, LO, Posterior Fusiform, pFuS) and body-selective (Extrastriate Body Area, EBA, Fusiform Body Area, FBA) areas. Using population receptive field (pRF) modeling, we obtain an estimate of spatial tuning at the single-voxel level, allowing us to investigate how spatial tuning relates to the strength of category-selectivity. With these data in hand, we critically evaluate two ways in which retinal sampling effects have previously been suggested to relate to category-selectivity.

First, we ask to what extent retinotopic biases can explain the presence of multiple different regions for the *same category* across *different* cortical surfaces. We (Groen et al., 2021; Kravitz et al., 2013; Silson et al., 2015) and others (Arcaro & Livingstone, 2017, 2021) earlier proposed that quadrant biases, inherited from splits in dorsal and ventral maps in early visual cortex (V1-V3), may explain why category-selective regions duplicate across lOTC and vOTC. We previously tested this retinotopic prediction for two scene-selective regions, one in lOTC, the OPA (Dilks et al., 2013) and one in vOTC, the PPA (Epstein & Kanwisher, 1998), and demonstrated that these regions (on average) exhibit biases for different portions of visual space with OPA showing a contralateral lower visual field bias and PPA a contralateral upper visual field bias (Silson et al., 2015, 2016). We speculated that such retinotopic biases would extend throughout lOTC and vOTC and not be confined to the scene-selective regions, specifically.

Second, we ask to what extent retinotopic biases can explain the presence of regions selective for *different categories* within the *same* cortical surface. We specifically focus on the center-periphery organizational framework for vOTC, which suggests that different category-selective regions emerge along the medial-lateral axis of vOTC based on the eccentricity in which we typically encounter those stimuli (Hasson et al., 2002; Levy et al., 2001). Selectivity for faces in FFA is thought to overlap foveal representations whereas selectivity for scenes in PPA is thought to overlap peripheral ones (Hasson et al., 2002; Levy et al., 2001). We first characterized the relationship between selectivity and pRF properties in FFA and PPA, before considering face- and place selectivity within vOTC more broadly.

## Methods

### Participants

Thirty participants completed the fMRI experiment (21 females, mean age = 24.2 years). All participants had normal or corrected to normal vision and gave written informed consent. The National Institutes of Health Institutional Review Board approved the consent and protocol. This work was supported by the Intramural Research program of the National Institutes of Health – National Institute of Mental Health Clinical Study Protocol 93-M-0170, NCT00001360.

### fMRI scanning parameters

Participants were scanned on a 3.0T GE Sigma MRI scanner using a 32-channel head coil in the Clinical Research Center on the National Institutes of Health campus (Bethesda, MD). Across all participants, whole brain coverage was acquired. Slices were oriented axially, such that the most inferior slice was below the temporal lobe. All functional images were acquired using a BOLD-contrast sensitive standard EPI sequence (TE = 30 ms, TR = 2 s, flip-angle = 65 degrees, FOV = 192 mm, acquisition matrix = 64×64, resolution 3 × 3 × 3 mm, slice gap = 0.3 mm, 28 slices). A high-resolution T1 structural image was obtained for each participant (TE = 3.47 ms, repetition time = 2.53 s, TI = 900 ms, flip angle = 7°, 172 slices with 1 × 1 × 1 mm voxels).

### Visual Stimuli and Tasks

In separate runs, participants were presented with two types of stimuli aimed at mapping two different types of selectivity: 1) visual field selectivity, using population receptive field task and 2) visual category-selectivity, using a six-category localizer task. All participants completed six population receptive field mapping runs and six category-localizer runs. Stimuli were presented using PsychoPy software (Peirce, 2007) (RRID:SCR_006571) from a Macbook Pro laptop (Apple Systems, Cupertino, CA). The tasks were alternated every two runs.

### Population receptive field (pRF) mapping

During pRF mapping sessions a bar aperture traversed gradually through the visual field, whilst revealing randomly selected scene fragments from 90 possible scenes (**Figure 1A)**. During each 36 s sweep, the aperture took 18 evenly spaced steps every 2 s (1 TR) to traverse the entire screen. Across the 18 aperture positions all 90 possible scene images were displayed once. A total of eight sweeps were made during each run (four orientations, two directions). Specifically, the bar aperture progressed in the following order for all runs: Left to Right, Bottom Right to Top Left, Top to Bottom, Bottom Left to Top Right, Right to Left, Top Left to Bottom Right, Bottom to Top, and Top Right to Bottom Left. The bar stimuli covered a circular aperture (diameter = 20° of visual angle). Participants performed a color detection task at fixation, indicating via button press when the white fixation dot changed to red. Color fixation changes occurred semi-randomly, with approximately two-color changes per sweep (Silson et al., 2015).

### Six category functional localizer

During each run, color images from six stimulus categories (Scenes, Faces, Bodies, Buildings, Objects and Scrambled images) were presented at fixation (10 × 10° of visual angle) in 16 s blocks (20 images per block [300 ms per image, 500 ms blank]) (**Figure 1B)**. Each category was presented twice per run, with the order of presentation counterbalanced across participants and runs. Participants responded via a MRI compatible button box whenever the same image appeared sequentially.

### fMRI data processing

#### Preprocessing

All data were analyzed using the Analysis of Functional NeuroImages (AFNI) software package (Cox, 1996) (RRID:SCR_005927). All functions and programs were available in the AFNI binary version April 21, 2020. Before pRF and functional localizer analyses, all images for each participant were motion corrected to the first image of the first run (*3dVolreg*), after removal of the appropriate “dummy” volumes (eight) to allow stabilization of the magnetic field. Post motion-correction data were detrended (*3dDetrend)* and, in the case of the localizer data only, smoothed with a 5 mm full-width at half-maximum Gaussian kernel (*3dMerge*).

### Population receptive field modeling

The population receptive field mapping runs were analyzed by fitting a pRF model (Dumoulin & Wandell, 2008) to the fMRI time courses of each voxel. Detailed description of the pRF model implemented in AFNI is provided elsewhere (Silson et al., 2015). Briefly, given the position of the stimulus in the visual field at every time point, the model estimates the pRF parameters that yield the best fit to the responses of a given voxel: pRF center location (x, y), and size (diameter of the pRF). Both Simplex and Powell optimization algorithms are used simultaneously to find the best time-series/parameter sets (x, y, size) by minimizing the least-squares error of the predicted time-series with the acquired time-series for each voxel.

### Six category functional localizer

Analyses were conducted using a general linear model approach using the AFNI programs *3dDeconvolve* and *3dREMLfit*. The data at each time point were treated as the sum of all effects thought to be present at that time point and the time series was compared against a Generalized Least Square (GLSQ) model fit with REML estimation of the temporal autocorrelation structure. Specifically, a response model was built by convolving a standard gamma function with a 16 s square wave for each condition and compared against the activation time courses using GLSQ regression. Motion parameters and four polynomials accounting for slow drifts were included as regressors of no interest. To derive the response magnitude per condition, *t*-tests were performed between the condition-specific beta estimates (normalized by the grand mean of each voxel for each run) and baseline.

### Sampling of data to the cortical surface

In each participant, the analyzed functional data were projected onto surface reconstructions (*3dvol2surf*) of each individual participant’s hemispheres derived from the Freesurfer4 autorecon script (http://surfer.nmr.mgh.harvard.edu/) using the Surface Mapping with AFNI (SUMA) software (Saad & Reynolds, 2012).

### Region of Interest (ROI) definitions

In each participant and hemisphere, ROIs were defined based on specific statistical contrasts (p<0.0001, uncorrected in each case). The number of participants in which an ROI was defined in each hemisphere were as follows: Scene-selective regions (Scenes > Faces), OPA (lh=30, rh=28), PPA (lh=29, rh=28). Face-selective regions (Faces > Scenes), OFA (lh=15, rh=16), FFA (lh=23, rh=27). Object-selective regions (Objects > Scrambled objects), LO (lh=15, rh=23), PFuS (lh=17, rh=14). Body-selective regions (Bodies > Objects), EBA (lh=23, rh=14), FBA (lh=25, rh=18).

### ROI analysis

ROIs were defined on the surface reconstructions using the interactive ROI drawing tool in SUMA and converted to a 1D index of node indices per ROI (*ROI2dataset*), which was subsequently used to extract both the pRF parameters (x, y, pRF size, R^2^) from the pRF modeling data and the selectivity profiles (t-value of each category versus baseline) from the functional localizer data (*ConvertDset*). The extracted data were then imported into Matlab (Version R2019B) for further analysis. Vertices were unique to each ROI (e.g. OPA vertices did not overlap those from OFA, EBA or LO).

### Visual field coverage and visual field biases

The visual field coverage plots represent the group average sensitivity of each region of interest (ROI) to different positions in the visual field. To compute these, individual participant visual field coverage plots were first derived. These plots combine the best Gaussian population receptive field model for each suprathreshold voxel (R^2^ > 0.2) within an ROI. Here, a max operator was used that reflects, at each point in the visual field, the maximum pRF value (i.e., height of the fitted Gaussian) from all pRFs within the ROI (Winawer et al., 2010). To compute visual field biases in individual participants and ROIs, we calculated the mean pRF value in the contralateral upper minus the contralateral lower visual field.

### Statistical analyses

Statistics were calculated using the R Studio package (version 1.3). For each category, we predicted an interaction between the lateral or ventral position of the ROI (i.e. Surface) and a preference for the upper visual field or lower visual field (i.e. Bias). For our analyses, we employed linear mixed modeling in order to assess the significance of Surface by Bias interactions, which are better able to account for missing data points than the less flexible repeated measures ANOVA.

## Results

### Visual field coverage

For each hemisphere and ROI, group-average visual field coverage plots were computed **(Figure 2 A-D)**. These plots represent a schematic visualization of the sensitivity of each ROI to different positions in the visual field and reveal three general patterns. First, each ROI exhibits a clear bias for the contralateral visual field, reflecting the privileged input of contralateral visual information - a hallmark of early retinotopically defined regions. Second, visual inspection reveals that in the majority of cases, lOTC ROIs (top row of each panel) exhibit a clear bias for the contralateral lower visual field, whereas vOTC ROIs (bottom row of each panel) exhibit either the opposite bias for the contralateral upper visual field or no clear quadrant bias. Third, there are qualitative differences in the degree of foveal versus peripheral sampling across ROIs with different category preferences. For instance, face-selective regions are most sensitive to foveal positions (**Figure 2B)**, followed closely by object-selective regions, whereas scene-selective regions show more pronounced peripheral positions (**Figure 2A)** with body-selective regions sensitive to more intermediate visual field positions (**Figure 2C-D)**.

**Figure 2.**
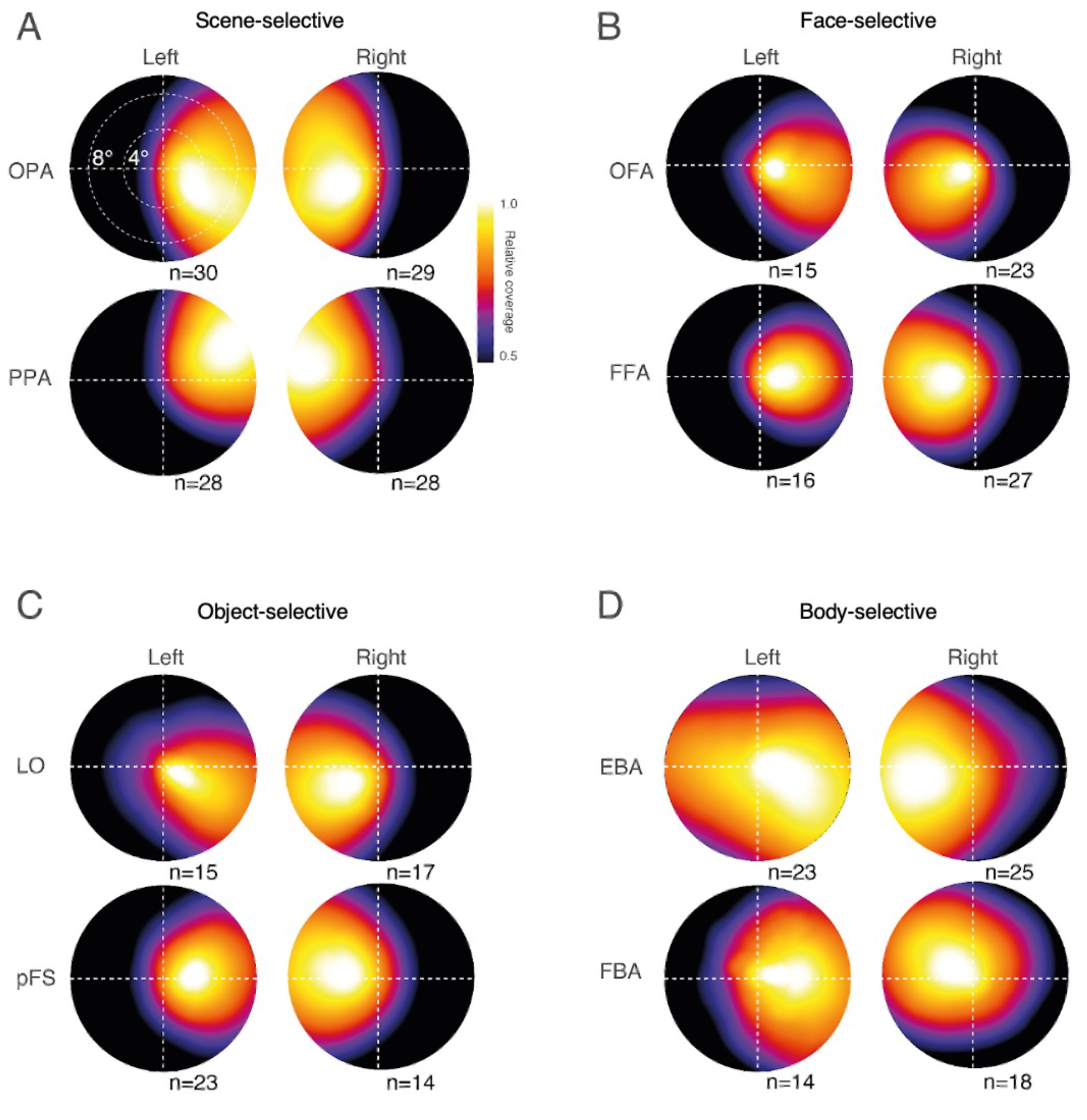
Visual field coverage. **A**, Group-averaged visual field coverage plots (10-10 dva) are shown for OPA (top row) and PPA (bottom row) for both hemispheres. The number of participants is inset. **B**, Same as A but for OFA (top row) and FFA (bottom row). **C**, Same as A but for LO (top row) and pFS (bottom row). **D**, Same as A but for EBA (top row) and FBA (bottom row). In each case a clear bias for the contralateral visual field is present. Despite subtle variation in the extent of the elevation bias across category-selective pairs, a consistent pattern remains evident. Lateral regions (top rows in A-D) all show a bias for the lower visual field, whereas ventral regions (bottom rows) either show the opposite bias for the upper visual field or no clear quadrant preferences.

### Within-category upper versus lower visual field biases

To quantify the presence of upper versus lower visual field biases, we performed the following analyses. For each participant and ROI, we calculated the average pRF value in the contralateral upper visual field and contralateral lower visual field, respectively. The resulting visual field bias indices were fitted with a linear mixed effects model. The model comprised three fixed effects, one for each within-subject factor: Hemisphere (Left, Right), Surface (Lateral, Ventral), and Bias (UVF, LVF), with participant added as a random effect. In each case, the presence of a significant Surface by Bias interaction was predicted, with an anticipated stronger LVF bias in lateral ROIs, but a stronger UVF bias in their ventral counterparts. These data are shown in **Figure 3 A-D (**averaged across hemispheres).

**Figure 3.**
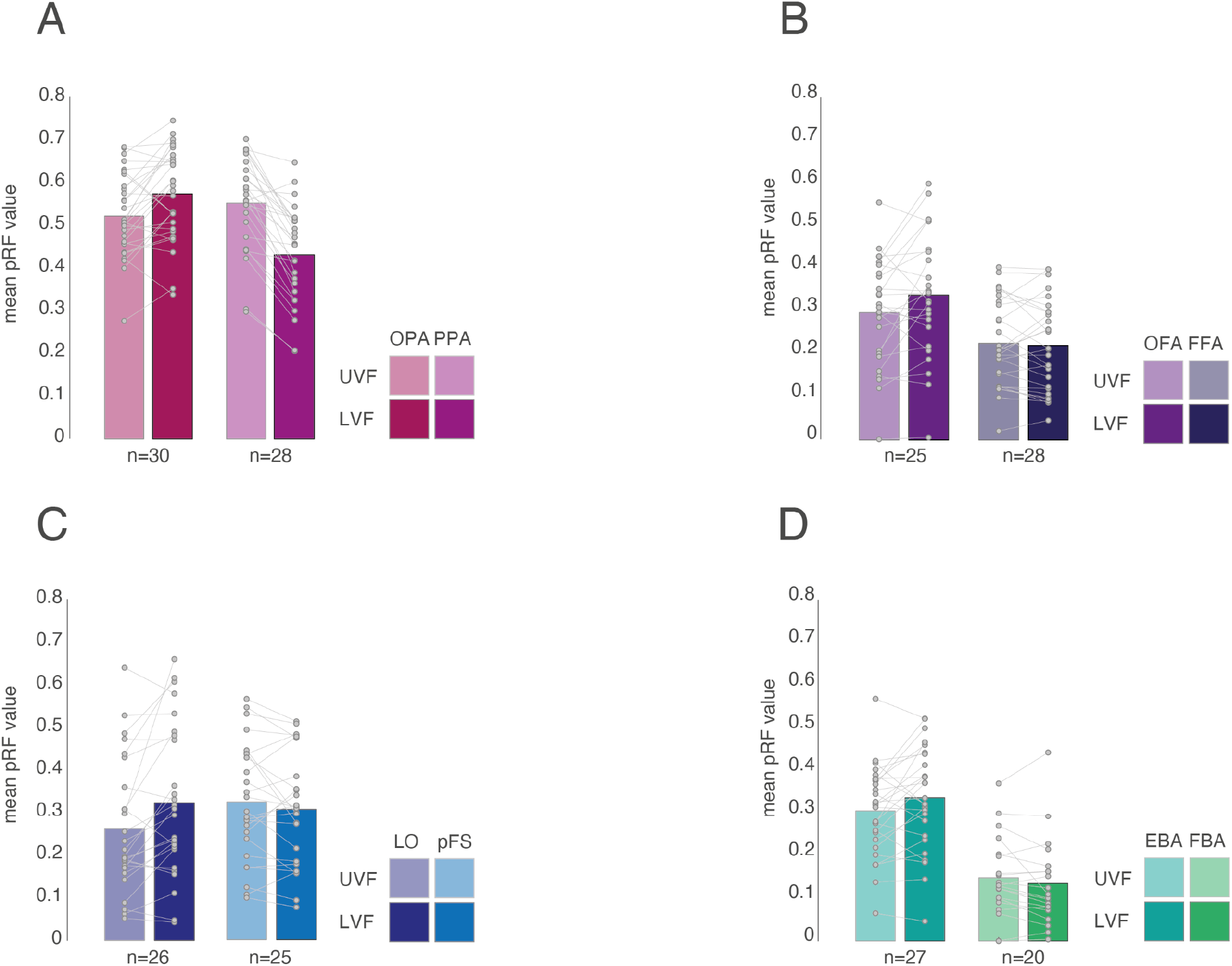
Visual field biases of category-selective pairs across lOTC and vOTC averaged across hemispheres. **A**, Bars represent the group-average pRF value in the contralateral upper visual field (faded bars) and contralateral lower visual field (solid bars) in OPA and PPA. In this and subsequent plots (B-D) individual participant data points are shown and linked for each ROI and the number of participants included is inset below. OPA shows a bias for the lower visual field, with PPA showing an upper visual field bias. **B**, As in A, but for OFA and FFA. OFA shows a bias for the lower visual field, but FFA shows no clear bias. **C**, As in A, but for LO and pFS. LO shows a lower visual field bias, with an upper field bias in pFuS. **D**, As in A, but for EBA and FBA. EBA shows a lower visual field bias with no clear bias in FBA.

#### Scene-selective regions (OPA vs PPA)

The main effects of Hemisphere (F(1, 28)=19.22, *p*=0.0001), Surface (F(1, 28)=80.59, *p*=9.85^-10^) and Bias (F(1, 28)=5.79, *p*=0.02) were significant, reflecting on average larger pRF values in the right than left hemisphere, on the lateral than ventral surface and for the upper over lower visual field, respectively. Importantly these were qualified by a significant Surface by Bias interaction (F(1, 112)=76.38, *p*=2.463^-14^), which reflects on average the larger LVF bias in OPA but larger UVF bias in PPA. All other interactions were not significant (*p*>0.05, in all cases).

#### Face-selective regions (OFA vs FFA)

The main effects of Hemisphere (F(1, 24.09)=5.89, *p*=0.02) and Surface (F(1, 28.15)=23.62, *p*=3.99^-5^) were significant, reflecting on average larger pRF values in the right than left hemisphere and on the lateral than ventral surface. The main effect of Bias (F(1, 28.34)=1.99, *p*=0.16) was not significant. Again, a significant Surface by Bias interaction (F(1, 97.52)=7.95, *p*=0.005) was observed. Unlike that observed for scene-selective regions, this interaction appears largely driven by the larger LVF bias in OFA as the UVF bias in FFA is marginal. All other interactions were not significant (*p*>0.05, in all cases).

#### Object-selective regions (LO vs pFS)

The main effects of Hemisphere (F(1, 23.08)=14.55, *p*=0.0008), Surface (F(1, 23.72)=7.76, *p*=0.01) and Bias (F(1, 111.37)=21.95, *p*=7.96^-6^) were significant, reflecting on average larger pRF values in the right than left hemisphere, on the lateral than ventral surface and in the LVF than UVF, respectively. Importantly and consistently, these main effects were qualified by a significant Surface by Bias interaction (F(1, 111.37)=25.50, *p*=1.73^-6^). The interaction reflects the larger LVF bias in LO, but larger UVF bias in pFS. All other interactions were not significant (*p*>0.05, in all cases).

#### Body-selective regions (EBA vs FBA)

Neither the main effects of Hemisphere (F(1, 24.34)=2.92, *p*=0.10) nor Bias (F(1, 24.99)=0.63, *p*=0.43) were significant, but the main effect of Surface (F(1, 21.00)=83.81, *p*=8.88^-9^) was, reflecting on average larger pRF values on the lateral than ventral surface. All interactions were not significant (*p*>0.05, in all cases). It is worth noting that the lack of interaction here might reflect the sparsity of FBA ROIs relative to EBA (22 fewer FBAs in the LMM).

Our first prediction tested whether consistent retinotopic biases for the lower and upper visual field could explain the paired selectivity across categories observed on the lateral and ventral surfaces of OTC. These results show that a significant Surface by Bias interaction was present in scene, face, object but not body-selective regions. These interactions reflect the fact that the locus of the visual field representation shifts from the contralateral LVF to the contralateral UVF as one moves from lOTC to vOTC. It is noteworthy, however, that although significant, the magnitude of this interaction differs between category-selective pairs.

### Peak eccentricity representations between categories

Our second prediction concerned whether consistent retinotopic biases could explain the presence of multiple category-selective regions within the lOTC and vOTC, respectively. First, we investigate this question within the lOTC and vOTC separately, before focusing on vOTC more directly.

In addition to upper and lower visual field biases within category-selective regions, Figure 2 also highlights differences in the eccentricity sampling between ROIs selective for different categories. To assess this formally, we first calculated, in each participant, an eccentricity vector that passed through the peak (i.e., hotspot) of each ROIs visual field coverage map before taking the median of this vector and averaging these values across hemispheres. Next, these averaged values were entered into a LMM with factors Surface (Lateral, Ventral) and Category (Scenes, Faces, Objects & Bodies) as fixed effects. As above, participant was modeled as a random effect. The main effect of Category (F(3, 175)=15.33, p=6.61-9) was significant, as was the main effect of Surface (F(3, 175)=6.612-9, p=0.003). The Surface by Category interaction (F(3, 174)=5.63, p=0.001) was significant, which is likely driven by the fact eccentricity values increased from lOTC to vOTC for all categories except bodies. Post-hoc pairwise comparisons (Bonferroni corrected) revealed only three significant cases: Scenes vs Faces (t(174)=4.99, p<0.0001), Scenes vs Objects (t(176)=4.74, p<0.0001) and Scenes vs Bodies (t(177)=6.12, p<0.0001). These data are shown in **Figure 4** (averaged across hemispheres).

**Figure 4.**
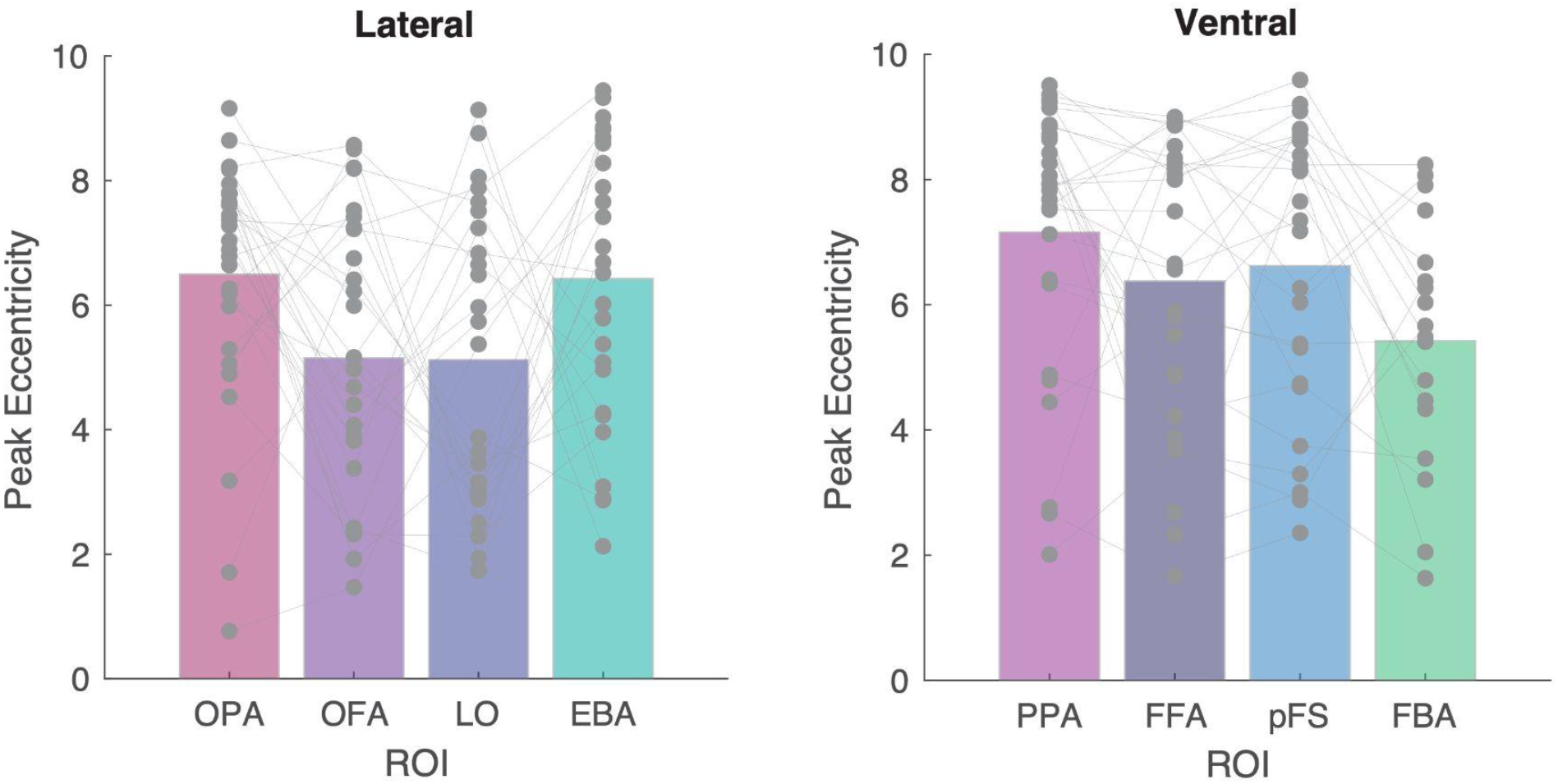
Peak eccentricity values between categories. **A**, bars represent the mean peak of eccentricity within each ROI on the lateral surface. Individual participant data points are plotted and linked across ROIs. **B**, same as A but for ROIs on the ventral surface.

**Figure 5** depicts the pRF center locations from FFA and PPA color-coded by their selectivity in both hemispheres pooled across all included participants (n=21). Several important patterns are evident within these data. First, although FFA pRFs are clustered at the fovea, PPA also contains substantial numbers of pRFs at the fovea. Second, both ROIs show pRFs extending peripherally into both the lower and upper visual field. Again, visual comparison reveals that these pRFs largely occupy similar visual field positions, but extend further into the periphery within PPA. Third, both FFA and PPA contain strongly selective pRFs (highly saturated colors) distributed throughout the visual field. In FFA, highly selective pRFs appear more frequently at the fovea but they also appear at peripheral visual field locations. In PPA, the highly selective pRFs appear to be more evenly distributed throughout the visual field. To quantify the degree of overlap in visual field representation between FFA and PPA we calculated, in each participant and hemisphere, the proportion of visual field coverage that is shared between FFA and PPA as a function of the total visual field represented by both FFA and PPA. This analysis revealed that FFA and PPA share ∼54% and ∼57% of their combined visual field coverage in the left and right hemispheres, respectively. Moreover, the vast majority of the remaining visual field coverage is unique to the PPA (lh: ∼33%; rh:∼30%) with unique FFA coverage representing only a relatively small percentage (lh: ∼13%; rh: ∼13%).

**Figure 5.**
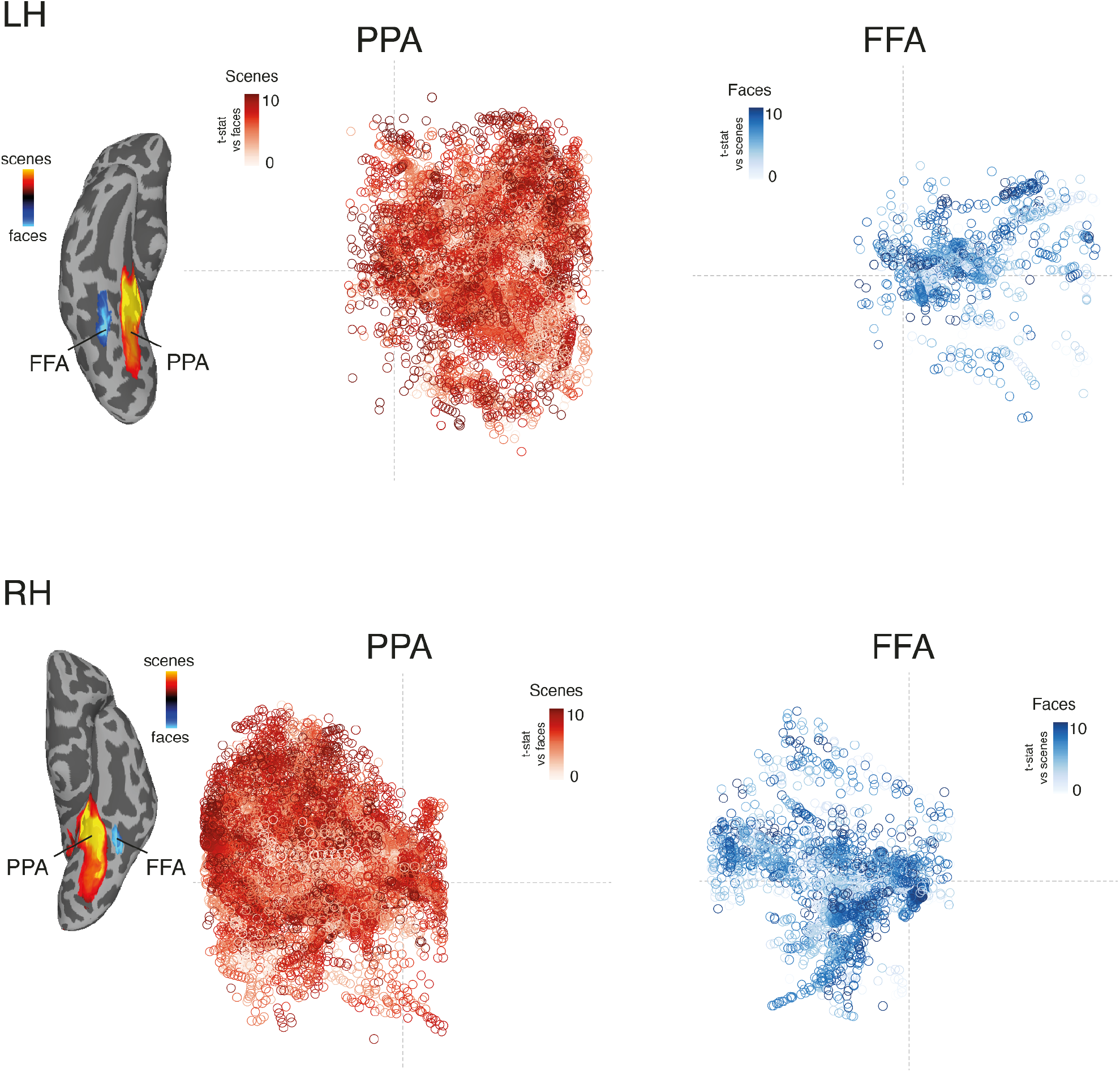
pRF center and selectivity in FFA/PPA. **Top**, Group-average ROIs for FFA (blue) and PPA (red) are inset. Overlaid onto the visual field are the centers of all suprathreshold (R2 >0.2) pRFs in PPA (red) and FFA (blue) color-coded by their corresponding selectivity (given by the *t*-value scenes v faces). Both PPA and FFA show prominent coverage of the fovea but also pRFs that extend throughout the visual field occupying very similar visual field positions. Both regions also show highly selective pRFs distributed throughout the visual field. **Bottom**, same as A but for PPA and FFA in the right hemisphere.

### Relationship between category-selectivity and pRF properties

Our analyses of visual field biases revealed both similarities and differences between category-selective regions. For example, all regions show robust coverage of the fovea, but whereas the peak of representation remains largely foveal in face- and object-selective regions, this peak is peripherally shifted in scene- and to a lesser extent body-selective areas. Further, regions in lOTC are collectively biased for the lower visual field with largely the opposite bias reflected in vOTC. Together, these data suggest that there is no simple unifying relationship between retinotopy and selectivity.

To explore this further, we next utilized the richness of our data set and focused on the center-periphery framework for vOTC, which suggests that it is governed according to an eccentricity gradient from foveal more laterally (i.e. FFA) to peripheral more medially (i.e. PPA) (Hasson et al., 2002; Levy et al., 2001). This center-periphery organization is thought to interact with visual experience to constrain the emergence of category-selectivity (Arcaro & Livingstone, 2017, 2021; Gomez et al., 2019). A strict interpretation of this framework suggests that there should be little to no overlap in visual field representation between lateral and medial areas of vOTC. Here, we tested this directly by characterizing pRFs in FFA and PPA in both hemispheres.

Having established that FFA and PPA share more visual field representation than commonly anticipated, we next explored the relationship between selectivity and eccentricity more broadly across vOTC. In each participant, we divided pRFs into five equal sections according to eccentricity (0-2 dva, 2-4, 4-6, 6-8 & 8-10) and computed the distribution of selectivity (given by the *t*-value scenes v faces) in each section. These distributions were then averaged across participants. These data reveal intriguing patterns of selectivity as a function of eccentricity **(Figure 6)**. First, consistent with the general center-periphery account, the proportion of scene-selective pRFs increases with eccentricity, whereas the proportion of face-selective pRFs is largest in the most foveal sections (two left-most plots) and decreases with increasing eccentricity. Second, even the most foveal sections contain relatively high proportions of pRFs that are stable as a function of scene-selectivity. That is, even highly scene-selective pRFs can occupy very foveal positions, so scene-selective pRFs in vOTC are not sampling the periphery exclusively. Third, even though the proportion of scene-selective pRFs increases with eccentricity, it is not the case that the most scene-selective pRFs are in the periphery. Indeed, fairly similar proportions of highly scene-selective pRFs are evident across eccentricities, and are, if anything, higher in more parafoveal than peripheral sections.

**Figure 6.**
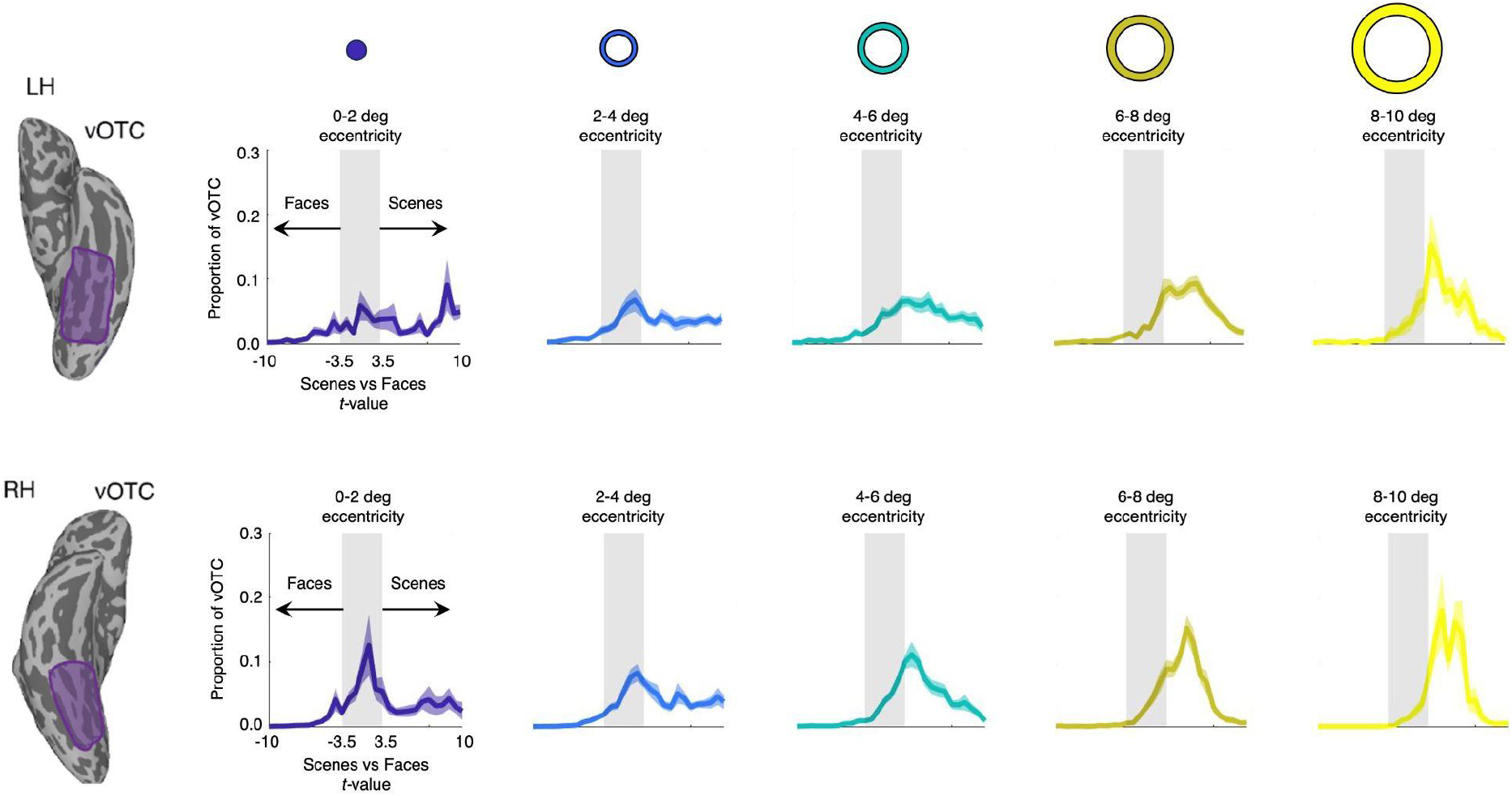
Selectivity distributions of foveal and peripheral pRFs within vOTC as a function of eccentricity in both hemispheres. The mean distributions of selectivity (plus sem) are plotted for five eccentricity sections: 0-2 dva (blue), 2-4 (cyan), 4-6 (green), 6-8 (mustard) and 8-10 (yellow). The shaded central area represents subthreshold selectivity values (*t >-3*.*5 or <3*.*5)*. The proportion of scene-selective pRFs increases with eccentricity, whereas the proportion of face-selective pRFs is largest in the most foveal sections (two left-most plots) and decreases with increasing eccentricity. Highly scene-selective pRFs are not restricted to peripheral locations but are present across eccentricities.

### Eccentricity and Scene-Selectivity in the Collateral Sulcus

Our vOTC analysis revealed that scene-selective pRFs are not restricted to the periphery, but are distributed fairly evenly across eccentricities. Further, visually comparing the spatial distribution of group-averaged scene-selectivity and eccentricity values suggests a gradient from peripheral to more-foveal along the posterior-anterior axis of vOTC, with the peak of scene-selectivity corresponding to relatively more foveal visual field representations **(Figure 8)**. Importantly, this gradient is distinct from the center-periphery gradient that runs along the medial-lateral axis of VTC.

This pattern prompted us to look more closely at the relationship between scene-selectivity and eccentricity within the collateral sulcus (CoS) specifically. Here, we first defined a series of line-ROIs spanning the length of the collateral sulcus along the posterior-anterior axis (see schematic inset, **Figure 9A**). Next, in each participant we sampled both the pRF eccentricity and scene-selectivity values along each line-ROI, before averaging across participants.

This analysis revealed a similar pattern in both hemispheres **(Figure 9)**. As a function of posterior-anterior location within the CoS, scene-selectivity initially increases, peaks and then decreases, whereas eccentricity begins peripherally (∼8 dva), becomes relatively more foveal, before returning to more peripheral. Strikingly, the peak in the scene-selectivity distribution corresponds spatially with the more-foveal trough of the eccentricity distribution, in both hemispheres, suggesting that the most scene-selective pRFs along the collateral sulcus are those at relatively more foveal eccentricities. These data suggest that a peripheral-foveal gradient exists along the posterior-anterior axis of the collateral sulcus that is independent of the commonly reported center-periphery gradient along the M-L axis of vOTC.

## Discussion

Here we leveraged detailed mapping of both visual field preferences and category-selectivity to investigate the systematic effects of retinotopic biases and category-selectivity on the organization of lOTC and vOTC, respectively. We had two main predictions: First, that different retinotopic biases for the upper and lower visual field could explain the paired selectivity found within lOTC and vOTC and second, that differences in eccentricity could explain differences between regions but within lOTC and vOTC separately.

Our data reveal both similarities and differences across regions and visual streams. We find that regions in lOTC exhibited clear contralateral lower visual field biases, whereas their corresponding vOTC regions exhibited either contralateral upper visual field biases or showed contralateral biases but with no clear quadrant preferences, confirming that lOTC and vOTC are indeed organized according to broad retinotopic gradients for different portions of the contralateral visual field (Arcaro & Livingstone, 2017, 2021; Groen et al., 2021; Kravitz et al., 2013; Silson et al., 2015). Importantly, despite these broad differences, similarities were also present across category-selective areas. Most consistent was that all regions showed robust foveal coverage with only the locus of the peak representation shifting between areas (e.g., from relatively more foveal in FFA to relatively more peripheral in PPA).

On the one hand, FFA and PPA showed the predicted differences in visual field biases in terms of eccentricity, consistent with the general tenant of the centre-periphery framework (Grill-Spector & Weiner, 2014; Hasson et al., 2002; Levy et al., 2001). Recent evidence also links viewing eccentricity with selective responses in experts over novices (Gomez et al., 2019) and the development of face-selectivity in the macaque (Arcaro et al., 2017; Arcaro & Livingstone, 2017, 2021) providing support for a strict eccentricity-dependent organization of category-selectivity in vOTC. On the other hand, FFA and PPA exhibited a surprisingly high degree of overlap in their pRF distributions, with ∼50% of the total visual field coverage being shared. Such a large degree of overlapping visual field coverage suggests the centre-periphery framework might benefit from a more nuanced interpretation. For instance, both FFA and PPA show prominent foveal coverage as well as pRFs that extend well into the periphery. What appears to differ between regions is the relative density and selectivity of pRFs in foveal and peripheral positions (**see Figure 5**). For example, FFA contains a relatively higher density of foveal pRFs, whereas PPA contains a relatively higher density of peripheral pRFs, but somewhat surprisingly, both regions contain strongly category-selective pRFs distributed throughout the visual field, inconsistent with a strict interpretation of the center-periphery framework driving face vs. place selectivity. Why might FFA and PPA have such shared visual field representations? One interpretation is that having different feature tuning at repeated positions in the visual field provides an efficient coding scheme for the visual cortex. Different preferences allow multiple visual features to be represented simultaneously, whilst the shared visual field representations between areas provides a means for rapid transfer of information. It remains to be seen what visual feature tuning preferences differentiate voxels in different category-selective regions that share a visual field representation. Important future work should seek to understand what visual features these regions are tuned to as a function of visual field position, particularly at shared/uniquely represented visual field representations (Groen et al., 2021).

**Figure 8.**
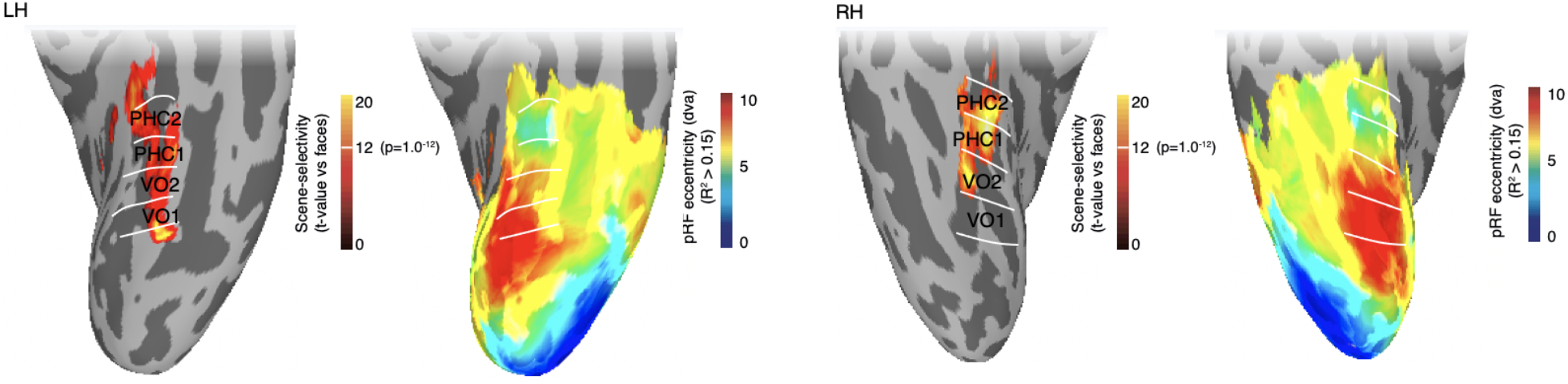
Scene-selectivity and eccentricity within vOTC. a ventral view of the right hemisphere is shown and enlarged to the right. Group-averaged scene-selectivity and eccentricity values are overlaid onto partially-inflated ventral surfaces of both the left-hemisphere (LH, left plot) and right-hemisphere (RH, right plot). Overlaid on hot-colors is the group-average scene-selectivity (thresholded at t>12, p=1.0-12). The borders of retinotopic maps VO1, VO2, PHC1 & PHC2 (taken from (Wang et al., 2015)) are overlaid in black. Group-average eccentricity map is overlaid onto the same view. Posterior portions of the collateral sulcus overlap with peripheral representations (red) and become relatively more foveal anteriorly.

**Figure 9.**
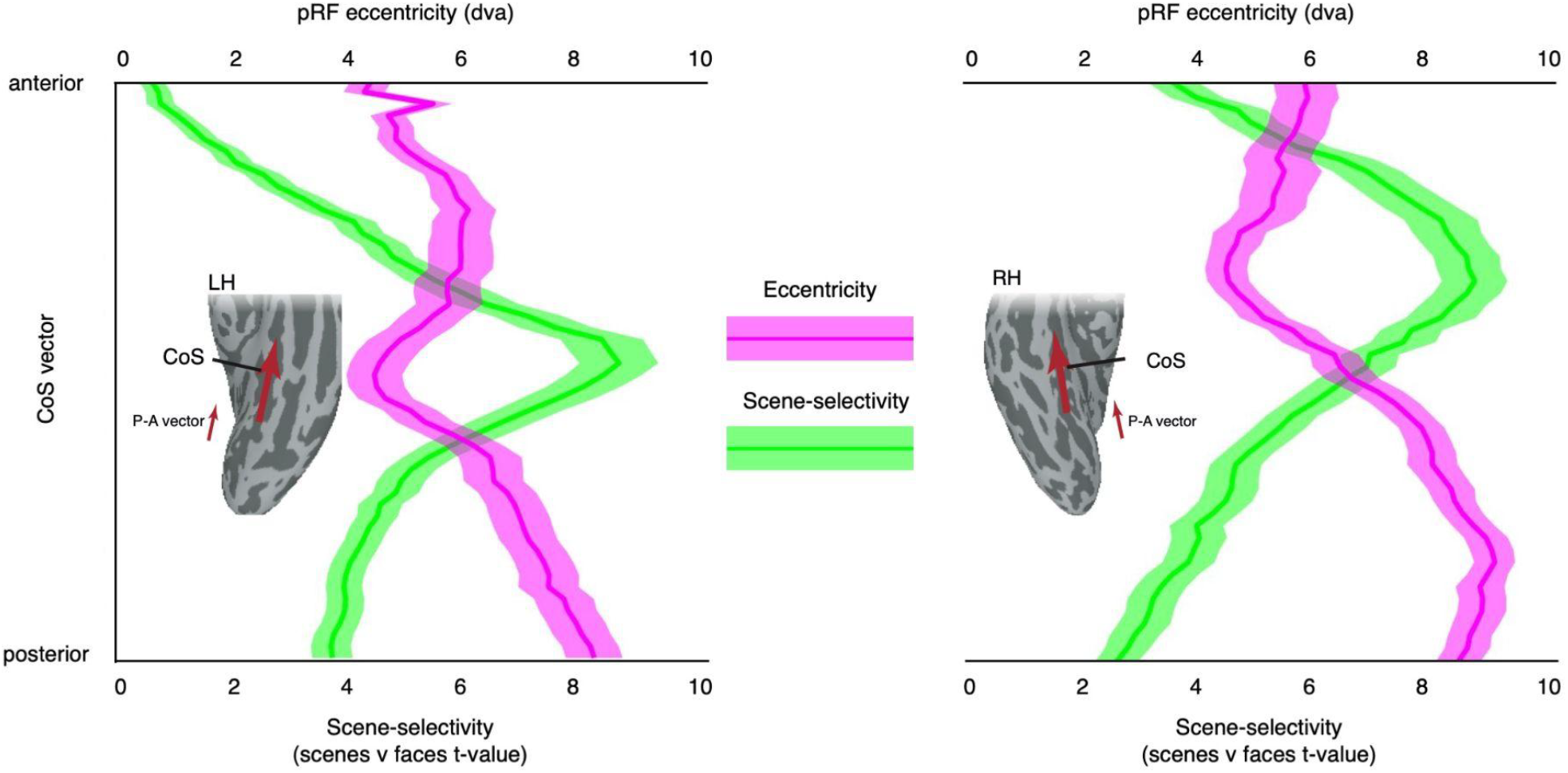
Eccentricity and scene-selectivity along the posterior-anterior axis of the collateral sulcus. Group-average mean (plus sem) scene-selectivity (green) and eccentricity (purple) distributions along the posterior-anterior axis are shown for the left hemisphere (LH, left plot) and right hemisphere (RH, right plot). The peak in the scene-selectivity distribution within the collateral sulcus corresponds spatially with the trough of the eccentricity distribution in both hemispheres.

Across vOTC more broadly, looking at face vs. place selectivity as a function of eccentricity, we again observe a pattern of results not fully consistent with the centre-periphery framework. While face-selective pRFs were more prominent within foveal over more peripheral eccentricities, the proportion of pRFs as a function of scene-selectivity were relatively stable within the most foveal eccentricities. Indeed, our data suggest that the most scene-selective pRFs are most prominent within parafoveal eccentricity and within the far periphery. These data highlight the complex relationship between eccentricity and selectivity, particularly in the case of responses to scenes. Finally, we investigated the relationship between eccentricity and scene-selectivity along the posterior-anterior axis of the CoS more closely. Here, we observe that alongside the well documented foveal-peripheral gradient along the L-M axis of vOTC (Grill-Spector & Weiner, 2014; Groen et al., 2021; Hasson et al., 2002; Levy et al., 2001), there is an additional and reverse peripheral-foveal gradient running along the posterior-anterior axis within the CoS. This gradient begins with peripheral representations posteriorly in the CoS, corresponding to the posterior portion of pPPA/PHC1 (Arcaro et al., 2009; Baldassano et al., 2013, 2016; Brewer et al., 2005). Consistent with the peripheral starting point of this gradient, both anatomical-(Beyh et al., 2022) and functional-connectivity data (Baldassano et al., 2016) indicate that pPPA receives biased input from regions of early visual cortex that represent the periphery. As a function of posterior-anterior location within the CoS, scene-selectivity initially increases, peaks and then decreases, whereas eccentricity begins peripherally (∼8 dva), becomes relatively more foveal, before returning to more peripheral. These data again highlight the complex relationship between eccentricity and selectivity in the context of scenes and suggest that the commonly accepted association of scene-selectivity and peripheral representations may be an oversimplification.

Overall, these data confirm the presence of robust visual field biases in category-selective areas and across the lateral and ventral processing streams. Importantly, however, different category-selective regions also share visual field representation. The considerable overlap for face vs. scene selective regions despite opposite selectivity profiles suggests different feature selectivities are computed at overlapping visual field positions. These data have important implications for theories about the origin of category-selectivity.

## Data and Code Availability

Data and code will be made available via the Open Science Framework upon acceptance.

## Author Contributions

EHS, IIAG and CIB jointly conceived of the project. EHS and IIAG collected the data. EHS & IIAG jointly analyzed the data and EHS, IIAG and CIB wrote the manuscript.

